# Low-dimensional representations of genome-scale metabolism

**DOI:** 10.1101/2024.05.31.596808

**Authors:** Samuel Cain, Charlotte Merzbacher, Diego A. Oyarzún

## Abstract

Cellular metabolism is a highly interconnected network with thousands of reactions that convert nutrients into the molecular building blocks of life. Metabolic connectivity varies greatly with cellular context and environmental conditions, and it remains a challenge to compare genome-scale metabolism across cell types because of the high dimensionality of the reaction flux space. Here, we employ self-supervised learning and genome-scale metabolic models to compress the flux space into low-dimensional representations that preserve structure across cell types. We trained variational autoencoders (VAEs) on large fluxomic data (*N* = 800, 000) sampled from patient-derived models for various cancer cell types. The VAE embeddings have an improved ability to distinguish cell types than the uncompressed fluxomic data, and sufficient predictive power to classify cell types with high accuracy. We tested the ability of these classifiers to assign cell type identities to unlabelled patient-derived metabolic models not employed during VAE training. We further employed the pre-trained VAE to embed another 38 cell types and trained multilabel classifiers that display promising generalization performance. Our approach distils the metabolic space into a semantically rich vector that can be used as a foundation for predictive modelling, clustering or comparing metabolic capabilities across organisms.

## I. INTRODUCTION

Genome-scale metabolic models (GEMs) are widely used to describe the connectivity of cellular metabolism. Dozens of algorithms have been developed to construct, curate and analyse these models for various applications^1^. In microbial systems, GEMs can be employed to understand the relation between metabolic phenotypes and growth conditions; in metabolic engineering, GEMs are widely used as tools to discover genetic modifications that maximize production of high-value compounds^2^. Other studies have developed GEMs for multicellular organisms^3^ as well as the human microbiome^4^. Most recently, GEMs have gained increased adoption to model human metabolism. A key development was the release of Recon3D, an all-encompassing model of human metabolism that can be instantiated to specific cell types with suitable omics’ data^5^. Common applications of human GEMs include the study of loss-of-function mutations in disease^6^ or the construction of patient-specific models for personalized medicine^7^.

A key challenge for the analysis of GEMs is their high dimensionality. For example, the most complete model for the *Escherichia coli* bacterium has more than 1,500 reactions, while human GEMs typically have well over 3,000 reactions. As a result, it is challenging to compare the properties of different GEMs and currently there is no unifying framework to describe high-dimensional GEMs across different organisms, cellular contexts or growth conditions. Earlier approaches for reducing the flux space employed mechanistic lumping of relevant pathways^8^, principal component analysis^9–11^, and various distance-based approaches to characterize differences across GEMs^12^.

Here, we employ self-supervised learning to build low-dimensional representations of GEMs. The approach is based on variational autoencoders (VAEs), a class of machine learning architectures widely employed in image processing, machine translation, and generative modelling^13^. We trained VAEs on a large corpus of fluxomic data generated via random sampling of GEMs derived from transcriptomic data of cancer patients spanning multiple tissues and cell types^14,15^. The trained models can be employed to compress flux samples into in a latent space of substantially lower dimension. Through clustering analysis and multilabel classification, we show that the VAEs embeddings can accurately reconstruct a high-dimensional flux space and, moreover, recover cell type identity with better accuracy than the fluxomic data employed for training. To the best of our knowledge, this is the first application of self-supervised learning for dimensionality reduction of GEMs. The approach offers potential as a novel strategy to characterize the metabolic capabilities of organisms, and contributes to the growing number of methods that combine machine learning with genome-scale metabolic modelling^11,16^.

## II. BACKGROUND

### A. Genome-scale metabolic modelling

A genome-scale metabolic model (GEM) describes cellular metabolism via the following relations:

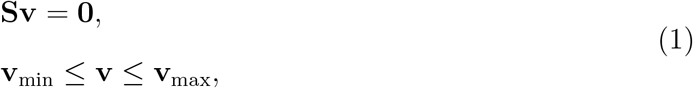

where **S** is an *m × n* integer matrix that describes the stoichiometry of a metabolic network with *m* metabolites and *n* reactions^17^, and **v** is an *n*-dimensional real vector of reaction fluxes. The *s*_*ij*_ entry of **S** describes the involvement of the *i*^th^ metabolite in the *j*^th^ reaction, with positive and negative values indicating metabolite production and consumption, respectively. The inequalities in (1) constrain the network fluxes to be within physiological bounds **v**_min_ and **v**_max_. A key aspect of this formulation is that GEMs can be instanced to specific cell types using proteomic or transcriptomic data to define the bounds **v**_min_ and **v**_max_. For example, reactions that are not present in a particular cell type can be removed from the model by zeroing out its corresponding bounds in (1), while reactions that are upregulated can be modelled through suitable scaling of their bounds.

From a geometric standpoint, the model in (1) defines a flux cone that contains all feasible metabolic states (Figure 1A). The flux cone is extremely high-dimensional, with typical human GEMs have in the order of *n* = 4, 000 to *n* = 6, 000 fluxes. This makes the characterization of its geometry particularly challenging. A number of approaches have been developed that exploit various properties of the flux cone, including extreme pathway analysis, elementary flux modes and others^1^. Among these approaches, flux sampling is a versatile framework that aims at characterizing the shape and size of the cone defined by (1). This is normally done using random walks suitably constrained to respect the geometry of the cone; the resulting collection of sampled flux vectors is typically referred to as *fluxomic* data.

**FIG. 1.**
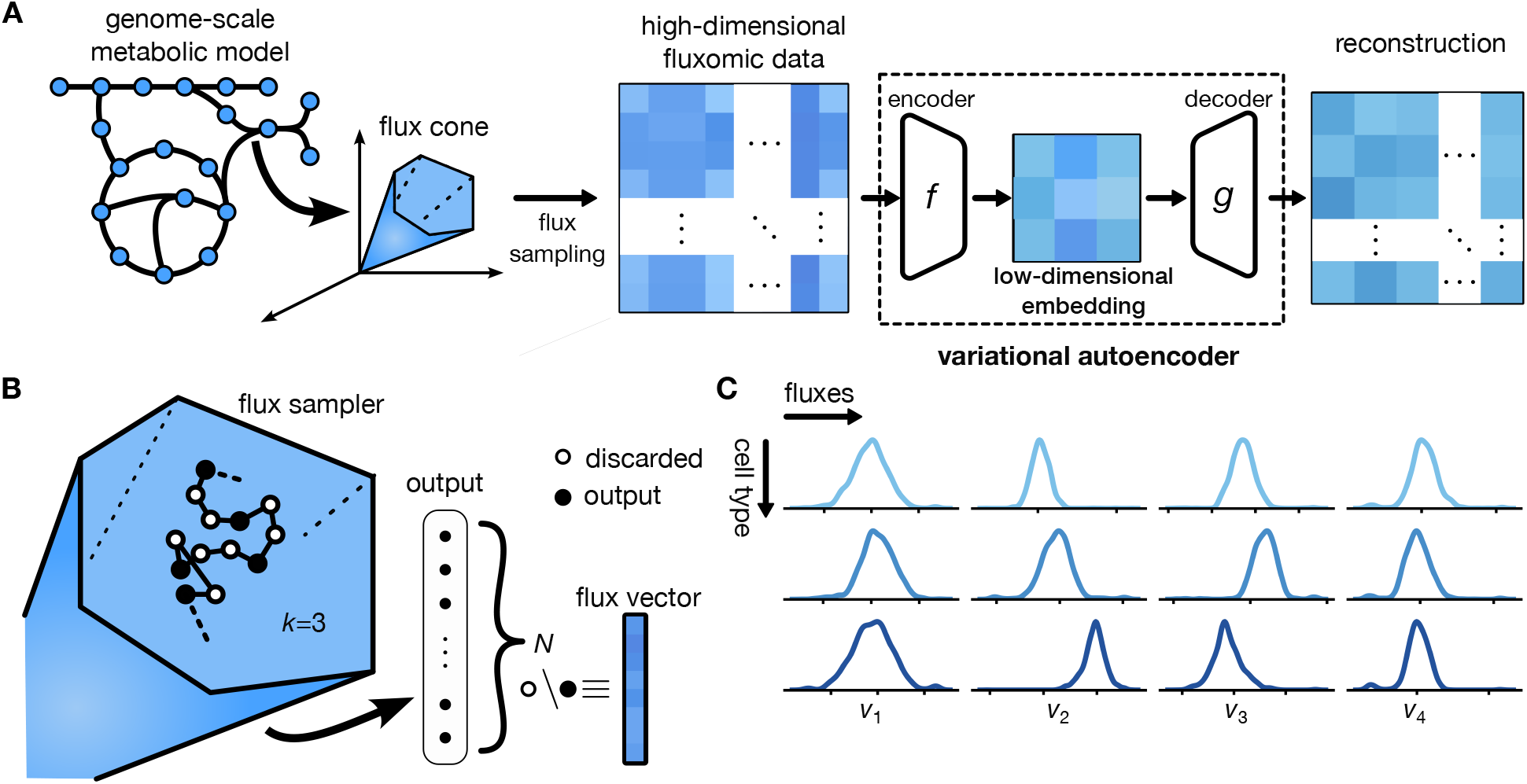
Pipeline for data generation and training of variational autoencoders (VAE) from fluxomic data. (**A**) We first generate training data by sampling of the flux cone associated to a genome-scale model (GEM). Prior to training, we prune inactive reactions (i.e. those with zero flux in at least one GEM) to ensure the data is not biased by the distribution of zero fluxes across cell types. The VAE compresses a flux sample down to a low-dimensional embedding space using the encoder, while the decoder decompresses the original sample. After training, the VAE embedding can be used for clustering or supervised prediction tasks. (**B**) Schematic of hit-and-run flux sampling with `OptGpSampler`^15^. Samples are generated via a discrete time random walk and kept only every *k* steps; in our case we employed *k* = 250. (**C**) Representative flux distributions obtained by sampling for various GEMs. Kernel density estimates of flux samples illustrate the variety of distributions present in the fluxomic data. Different cell types have different flux distributions for the same reaction, and flux distributions can vary widely across reactions.

In this paper, to generate training data we made extensive use of flux sampling with an implementation of the `OptGpSampler` available in the CobraPy python package^15,18^. In a first warm-up phase, the algorithm generates a collection of feasible flux vectors, which are then employed to initialize a random walk that generates samples using a hit-and-run approach (Figure 1B). The algorithm biases the choice of direction choice to be in the least constrained directions allowing larger step sizes between neighbours in the walk. As illustrated in Figure 1C, for a single GEM the resulting fluxomic data is composed of *n* distributions with *N* samples each. In the case of `OptGpSampler`, a single sample is saved every *k* discarded samples, and thus *k* can be used for controlling the trade-off between runtime and sampling quality.

### B. Representation learning

In machine learning, feature or representation learning refers to a suite of techniques employed to discover informative descriptions of data that can be employed for downstream tasks such as clustering, dimensionality reduction or classification. A common architecture for this task are variational autoencoders (VAEs), whereby two neural networks, an encoder *f* and a decoder *g* (Figure 1A), are jointly trained in a self-supervised fashion. The encoder networ takes an input vector **v** and maps it into a low-dimensional latent (or *embedding*) vector **z**. The decoder network then reverses this mapping by producing a reconstructed vector **v**_*r*_ of the same dimensionality as the input. The coordinates of a sample in the embedding space can then be used for downstream supervised or unsupervised tasks.

In its most common formulation, VAE training involves a loss function of the form

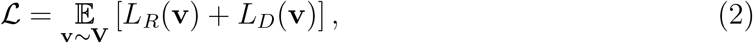

where *L*_*R*_ and *L*_*D*_ are the reconstruction and divergence loss, respectively, and the minimization is done over the expected loss across all samples **v** uniformly sampled from the training data **V**. The reconstruction loss is defined as:

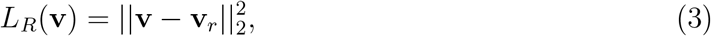

and its minimization ensures that the model accurately reconstruct the samples in terms of their euclidean distance. This ensures that the encoder defines an embedding mapping which preserves information needed for reconstruction. The divergence loss, on the other hand, is defined as:

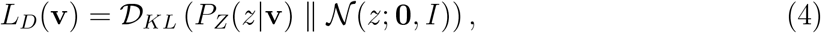

where *D_KL_* is the Kullback-Leibler (KL) divergence between the distribution over the embedding space and a target Gaussian with zero mean and unit-variance *N*(*z*; 0, *I*). The term *P*_*z*_(*z*|**v**) is the distribution over the embedding space for an input vector **v** as defined by the encoder. Minimizing the divergence loss ensures that the distribution of samples in the embedding space approaches a Gaussian, implicitly ensuring interpolations between input embeddings are represented by the neural network.

## III. MODEL TRAINING AND TUNING

### A. Data generation and aggregation

To generate the training data we ran `OptGpSampler` on 40 GEMs for various cancer types derived from patient transcriptomic data^14^. Through grid search we determined a step size *k* = 250 as a suitable compromise between compute resources and the quality of the resulting flux distributions. Since the sampler does not satisfy the Markov property, it is not guaranteed to converge to the uniform distribution^15^. Thus, to avoid confounders caused by specific sampler runs, we ran `OptGpSampler` 4 times for each GEM with *N* = 5, 000 samples per run using 32 CPU cores in parallel, totalling 20, 000 samples per cell type. To aggregate the fluxomic data of the 40 cell types into a single dataset, we discarded all reactions that were inactive (i.e. have zero flux) in at least one GEM. In other words, the aggregated fluxomic dataset contains only reactions that are active in all considered GEMs, which we term *core* reactions. This data pruning step was needed to ensure that all samples have the same dimension and prevent the VAE from simply learning the pattern of zeroes in each GEM. Overall, the training data has 40*×*20, 000 = 800, 000 samples of dimension *d* = 1, 675. Prior to training, training all flux samples were normalized to have zero mean and unit variance. Test data was produced with the same approach using 2 sampling runs per GEM, totalling 400, 000 flux vectors for model testing.

### B. Model architecture and training

We employed feed-forward neural networks for both encoder and decoder. Since these networks must invert each other, we chose them to have similar complexity and thus the same number of hidden layers. Through grid search we found that 3 hidden-layers for each network gave a good balance between training efficiently and achieving high flexibility in the networks (not shown). The sizes of the encoder input and decoder output were fixed by the number of core reactions, in our case *d* = 1, 675, while the number of nodes in the hidden layers were calculated as a linear interpolation between the dimensions of the input and output layers. We employed ReLU activation functions for the hidden layers and a linear activation for the output layers. Dense connections were used between all layers. For training we performed mini-batch gradient descent using the Adam optimizer with parameters *β*_1_ = 0.9 and *β*_2_ = 0.999. The VAE showed stable training with a batch size of 256 and a learning rate of 1 *×* 10^*−*4^; a weight decay of 0.05 was used to regularize the networks.

To determine the optimal dimension of the embedding space (*D*_emb_), we trained VAEs with *D*_emb_ = {2, 8, 32, 128, 512} and qualitatively evaluated their performance using the t-distributed Stochastic Neighbour Embedding (t-SNE) algorithm for nonlinear dimensionality reduction^19^. As shown in Figure 2A, the similarity between t-SNE representations of the fluxomic data, low-dimensional embedding, and the reconstructed flux samples suggests a promising performance of the VAE strategy, even for dimensions as low as *D*_emb_ = 8. The structure of the data, as captured by the t-SNE representation, seems to be well preserved in all cases except *D*_emb_ = 2. For a more quantitative evaluation of the impact of the embedding dimension, we evaluated the minimal test loss (Figure 2B). Larger *D*_*emb*_ gives the network more dimensions to encode **v** and we observed a plateau and increase in the loss for the larger embedding sizes. This suggests the VAE is not able to identify extra features that can improve the reconstruction. We thus chose to proceed with an intermediate embedding size *D*_emb_ = 32 to balance a low training loss with the increased tractability of a low-dimensional latent space.

**FIG. 2.**
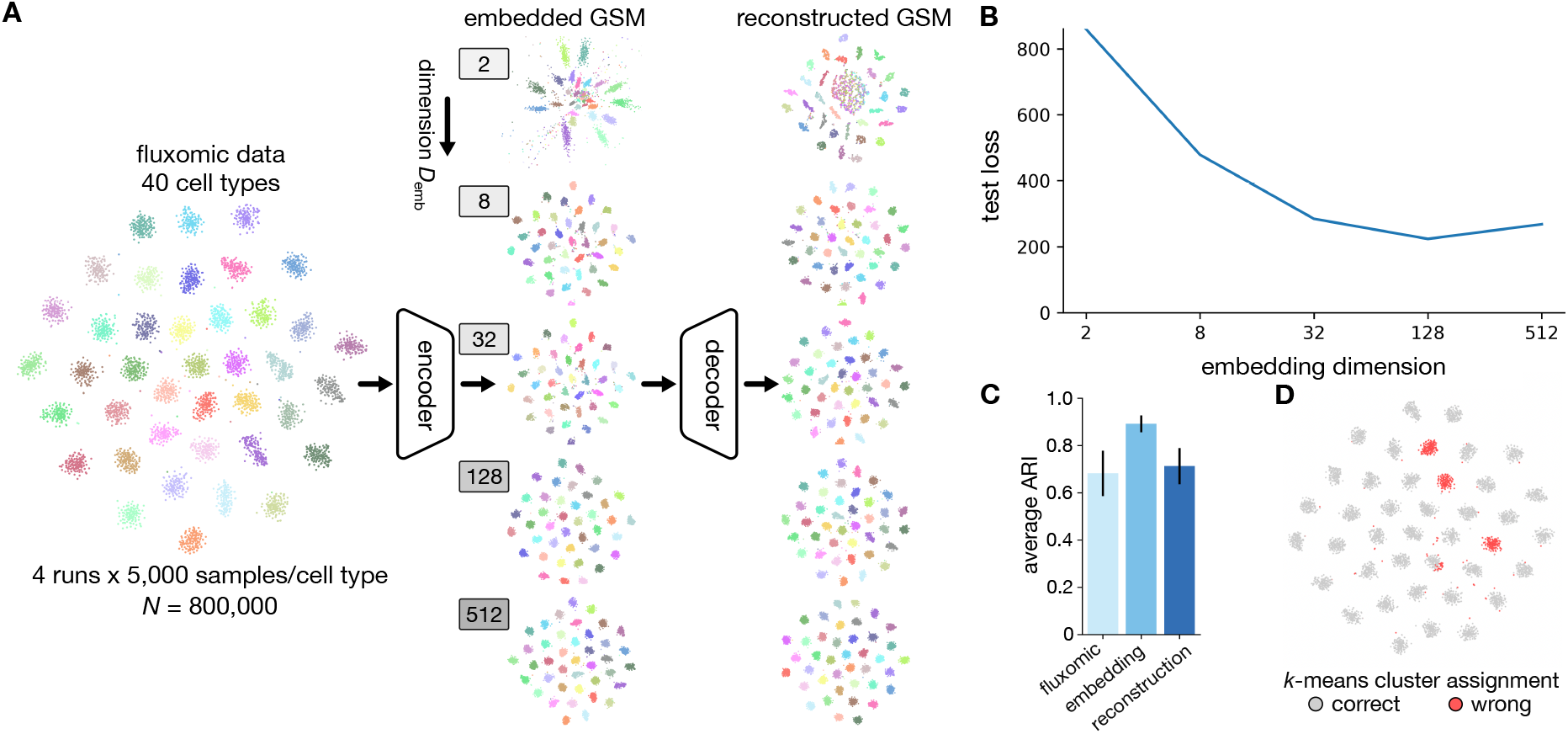
Performance of variational autoencoder trained on fluxomic data from various cell types. (**A**) We trained VAEs of increasing embedding dimension (*D*_emb_) on *N* = 800, 000 samples from 40 patient-derived GEMs for various cancer types; after pruning inactive reactions, the fluxomic data has *d* = 1, 675 dimensions. Shown are t-SNE visualizations of a sample (*N* = 8, 000; 200 samples per cell) from the test data withheld from training (left), compressed by the VAE embedding (middle) and reconstructed by the decoder neural network (right). The embedding and reconstructed flux samples appear to qualitatively preserve the structure of the data; colors correspond to the cell type were the raw fluxomic data was sampled from. The t-SNE visualizations were produced with `scikit-learn` in Python using perplexity of 30, early exaggeration of 12, and an adaptive learning rate. For *D*_emb_ = 2 we plot the embedding itself. (**B**) Minimum test loss achieved for embeddings of increasing dimension (*D*_emb_), computed on the withheld sampling runs employed as test data. (**C**) Similarity between *k*-means cluster assignment (*k* = 40) and the original cell type labels. Shown are the Adjusted Rand Index (ARI) scores between ground truth and raw fluxomic data, low-dimensional embedding, and the reconstructed fluxomic data. Bars denote average ARI *±* one standard deviation across 40 runs of the *k*-means clustering algorithm. (**D**) t-SNE visualization of misclustered samples in the embedding space with dimension *D*_emb_ = 32.

## IV. LEARNED REPRESENTATIONS PRESERVE STRUCTURE OF FLUXOMICS DATA

To evaluate the ability of the flux embeddings to preserve the structure of the metabolic space across cell types, we performed unsupervised and supervised learning with the learned embeddings as feature vectors. In both cases, the aim was to establish if the embeddings can recover the cell type identity of the fluxomic data.

### A. Clustering of learned flux embeddings

Although the t-SNE plots in Figure 2A suggest a correspondence between the structure of the fluxomics data and the low-dimensional embedding, nonlinear dimensionality reduction methods are prone to misrepresentations and visual artefacts. To quantitatively analyze the cluster structure of the embedded GEMs, we performed *k*-means clustering with *k* = 40 on the original fluxomic data (**v**), the learned embeddings (**z**), and the VAE reconstructed samples (**v**_*r*_). We compared the resulting cluster assignments with the 40 cell type labels using the Adjusted Rand Index (ARI) as a measure of similarity between clusterings; an ARI close to nil indicates little correspondence between cluster assignments, and an ARI of unity is achieved when two clusterings are identical.

The results in Figure 2C show that the learned embeddings provide the closest similarity the ground truth cell types, achieving a high average ARI of 0.89, while the *k*-means clusters of the original and reconstructed data only achieve an average ARI of 0.68 and 0.71, respectively. Moreover, we observed a smaller variance in ARI scores in the embedding space across *k*-means repeats with random initialization, which suggests that the VAE embedding can robustly preserve information on cell type. The improved performance of the *k*-means clustering on the embedding space could result from the divergence loss driving the embedding distributions towards isotropic Gaussians that match the *k*-means prior. As seen in Figure 2D, the samples wrongly assigned by *k*-means appear to be concentrated mostly in three cell types, while samples from the other 37 cell types are generally well clustered in the embedding space.

### B. Classification

To further explore the representative power of the embedding space, we trained 40-label classifiers using the VAE embeddings as feature vectors and several classification models: nearest centroid (NC), multi-layer perceptron (MLP) and random forest (RF). We trained the classifiers on the same data employed to train the VAE but embedded in the latent space with *D*_emb_ = 32; likewise, the classifiers were tested on the same sampler runs employed to test the VAE in Figure 2A. The results in Figure 3A show that the NC classifier achieved a near perfect accuracy above 99.95%, verifying that the VAE fits cells to regular clusters as suggested by *k*-means clustering. The MLP achieved a similar accuracy of 99.91% suggesting that the deeper architecture of the decoder has cell type available during the decoding process. The RF model with max-depth of 2 (RF-2 in Figure 3A) has the worst accuracy at 90.74%, although when the max-depth was increased to 4 (RF-4 in Figure 3A) the accuracy improved to 97.42%. We also observed that the accuracy of RF-2 varied widely between cell types, suggesting the VAE has learned to map some cell types to more isolated regions of the embedding space. The performance of the RF classifier can likely be improved with further hyperparameter optimization.

**FIG. 3.**
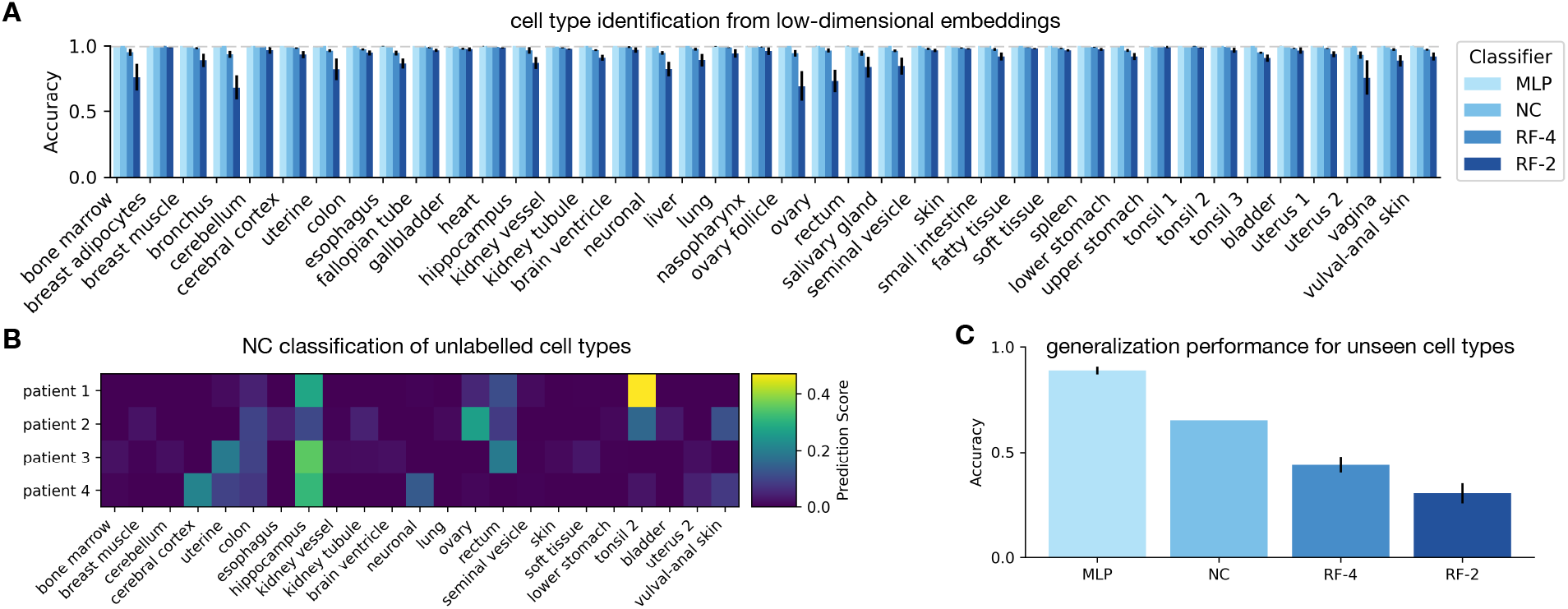
Performance of low-dimensional embeddings for cell type classification. (**A**) We trained 40-label classifiers on the embeddings produced by the VAE with *D*_emb_ = 32 and the ground truth cell type labels (Figure 2A). Four classification models were employed: nearest centroid (NC), multilayer perception (MLP) and random-forest with max-depth 2 (RF-2) or 4 (RF-4). The MLP had a single hidden layer with 100 nodes and ReLU activations. Each cell type was equally represented in the training data to avoid class imbalance. Accuracy scores were computed on the same test samples in Figure 2A (*N* = 8, 000; 200 samples per cell type); error bars denote one standard across 40 rounds of retraining of MLP and RF classifiers with different random seeds. (**B**) Predictions of NC classifier from panel A when queried with GEMs for liver cancer patients that were not used for VAE training; these GEMs were presented by Agren, R. et al as patients 2177, 2280, 2556 and 3196, respectively. We sampled each GEM and embedded them with the pre-trained VAE from Figure 2A with *D*_emb_ = 32; per-label prediction scores below 0.01 were excluded from the heatmap. (**C**) Classification performance on cell types not used in VAE training. We trained 38-label classifiers on the VAE embeddings of data sampled from additional 38 cell type GEMs that were not employed for VAE training^14^. The new training and test data were created with the same flux sampling parameters as the data for VAE training. Classifiers were trained as in panel A and tested on a sample (*N* = 7, 600; 200 per cell type) from the new test set.

To test the utility of the learned embeddings to assign cell types to unlabelled data, we employed the trained 40-label classifier from Figure 3A to predict cell type identity from new GEMs parameterized with transcriptomic data from liver cancer patients. Such patient-specific GEMs are particularly useful when examining the metabolic impact of various cancer mutations. We selected four liver cancer models from Agren, R. et al and embedded them using the VAE from Figure 2A with *D*_emb_ = 32. We queried the pre-trained NC classifier from Figure 3A for cell type predictions and obtained prediction scores across the 40 cell types (Figure 3B). These scores reflect how samples from unlabelled GEMs are distributed when embedded in the pre-trained VAE, and may provide a novel route to compare GEMs across cell types.

We finally tested the generalization performance of the learned embeddings and their ability to describe cell types not employed for VAE training. We constructed a new dataset by sampling 38 remaining GEMs presented by Agren, R. et al. We used the same strategy and sample size to construct this new fluxomic dataset from the GEMs unseen in VAE training. We embedded the new data using the pre-trained VAE from Figure 2A with *D*_emb_ = 32, and used the embeddings as feature vectors to train 38-label classifiers. As seen in Figure 3C, the accuracy of the NC model was 65.26% while the RF achieved an average of 30.46% and 43.91% for a max-depth of 2 and 4, respectively. The significantly lower accuracy of these classifiers suggests that the embeddings do not cluster into well separated groups. However, we found that the MLP achieved an excellent average accuracy of 88.82%, suggesting that the VAE has learnt general features that generalize across cell types.

## V. CONCLUSION

Genome-scale metabolic models (GEMs) are widely used to describe metabolic phenotypes across organisms and environmental conditions. At its core, a GEM defines a cone in a high-dimensional flux space. Several strategies have been developed to characterize the geometry of the flux cone, many of which involve random sampling or computation of geometric properties such as extreme pathways or elementary flux modes^1^. Recent years have witnessed a growing interest in the use of machine learning in tandem with GEMs^11,16^, particularly for improving phenotype predictions^20,21^, enrich the quality of metabolic models^22,23^, or interpreting gene expression data^24^.

In this paper, we proposed a machine learning approach to compress the flux cone into a low-dimensional latent space that preserves structure across cell types. Our results suggest that variational autoencoders can achieve substantial reductions in dimensionality, without compromising the representation power of the learned embedding. We tested the approach using a large fluxomic dataset and showed that cell type identity can be recovered from the embeddings with excellent accuracy. A salient property of our method is that the embeddings appear to be better clustered than than the fluxomic data employed for training. This suggests that the VAE strategy can increase the separability of the learned embedding and allow for easier reconstruction of cell type identity. Our computational experiments suggest that prediction based on the learned embeddings can generalize to cell types not employed for VAE training. Moreover, we demonstrated a proof-of-concept application to assign cell type identity to unlabelled metabolic models.

Further applications of our approach include the construction of metabolic distances between cell types, comparison across disease states and patient-specific GEMs, or using the learned embeddings for downstream prediction tasks. We envision this work as a new direction toward tractable representations of metabolism at the genome scale.

## ACKNOWLEDGEMENTS

CM and DAO were supported by the United Kingdom Research and Innovation (grant EP/S02431X/1, UKRI Centre for Doctoral Training in Biomedical AI).

## REFERENCES

1 N. E. Lewis, H. Nagarajan, and B. O. Palsson, “Constraining the metabolic genotype-phenotype relationship using a phylogeny of in silico methods,” Nat Rev Microbiol 10 (2012).

2 Liu, Di et al, “Dynamic metabolic control: towards precision engineering of metabolism,” J Ind Microbiol Biotechnol 45 (2018).

3 Gebauer, J. et al., “A genome-gcale database and reconstruction of Caenorhabditis elegans metabolism,” Cell Syst 2 (2016).

4 Heinken, A. et al., “Genome-scale metabolic reconstruction of 7,302 human microorganisms for personalized medicine,” Nat Biotechnol 41 (2023).

5 Brunk, E. et al., “Recon3d enables a three-dimensional view of gene variation in human metabolism,” Nat Biotechnol 36 (2018).

6 A. Mardinoglu and J. Nielsen, “New paradigms for metabolic modeling of human cells,” Curr Opin Biotechnol 34 (2015).

7 J. E. Lewis and M. L. Kemp, “Integration of machine learning and genome-scale metabolic modeling identifies multi-omics biomarkers for radiation resistance,” Nat Commun 12 (2021).

8 Ataman, M. et al., “redGEM: Systematic reduction and analysis of genome-scale metabolic reconstructions for development of consistent core metabolic models,” PLOS Computational Biology 13 (2017).

9 C. L. Barrett, M. J. Herrgard, and B. Palsson, “Decomposing complex reaction networks using random sampling, principal component analysis and basis rotation,” BMC Systems Biology 3, 30 (2009).

10 E. Yaneske and C. Angione, “The poly-omics of ageing through individual-based metabolic modelling,” BMC Bioinformatics 19 (2018).

11 Antonakoudis, A. et al., “The era of big data: Genome-scale modelling meets machine learning,” Comput Struct Biotechnol J 18 (2020).

12 A. Cabbia, P. A. J. Hilbers, and N. A. W. van Riel, “A Distance-Based Framework for the Characterization of Metabolic Heterogeneity in Large Sets of Genome-Scale Metabolic Models,” Patterns 1, 100080 (2020).

13 D. P. Kingma and M. Welling, “Auto-encoding variational bayes,” 1312.6114 [cs, stat] (2022).

14 Agren, R. et al., “Identification of anticancer drugs for hepatocellular carcinoma through personalized genome-scale metabolic modeling,” Mol Syst Biol 10 (2014).

15 W. Megchelenbrink, M. Huynen, and E. Marchiori, “OptGpSampler: an improved tool for uniformly sampling the solution-space of genome-scale metabolic networks,” PLoS ONE 9 (2014).

16 Zampieri, G. et al., “Machine and deep learning meet genome-scale metabolic modeling,” PLoS Comp Biol 15 (2019).

17 X. Fang, C. J. Lloyd, and B. O. Palsson, “Reconstructing organisms in silico: genome-scale models and their emerging applications,” Nat Rev Microbiol 18 (2020).

18 Ebrahim, A. et al., “Cobrapy: Constraints-based reconstruction and analysis for python,” BMC Syst Biol 7 (2013).

19 L. van der Maaten and G. Hinton, “Visualizing data using t-sne,” J Mach Learn Res 9 (2008).

20 Faure, L. et al., “A neural-mechanistic hybrid approach improving the predictive power of genome-scale metabolic models,” Nat Commun 14 (2023).

21 R. Hasibi, T. Michoel, and D. A. Oyarzún, “Integration of graph neural networks and genome-scale metabolic models for predicting gene essentiality,” NPJ Syst Biol Appl 10 (2024).

22 Li, F. et al., “Deep learning-based kcat prediction enables improved enzyme-constrained model reconstruction,” Nat Catal 5 (2022).

23 Pio, G. et al., “Integrating genome-scale metabolic modelling and transfer learning for human gene regulatory network reconstruction,” Bioinformatics 38 (2022).

24 Barsacchi, M. et al., “GEESE: Metabolically driven latent space learning for gene expression data,” bioRxiv bioRxiv: 10.1101/365643 (2018), https://www.biorxiv.org/content/early/2018/07/11/365643.full.pdf.

